# Conditional deletion of human STN1 leads to telomere dysfunction and telomerase-dependent genome instability and proliferation defects

**DOI:** 10.1101/2025.08.06.669015

**Authors:** Jaclyn S. Holbrooks, Colin A. Loveless, S. Donte’ Reed, Grayson H. Duvall, Carlan V. Romney, Madison B. Kircher, Jason A. Stewart

## Abstract

CTC1-STN1-TEN1 (CST) is a heterotrimeric, RPA-like complex that binds single-stranded DNA, stimulates DNA polymerase α-primase, and functions in several genome maintenance pathways, including telomere maintenance and DNA replication/repair. During telomere replication, CST prevents telomerase from overextending the G-rich single-stranded overhang (G-OH) and promotes fill-in of the C-rich strand by stimulating DNA polymerase α-primase. Previous work characterized the effects of CST loss by deleting CTC1 or TEN1. Interestingly, CTC1 knockout (KO) caused severe proliferation defects and telomeric damage signaling, whereas these phenotypes were absent following TEN1 KO. Molecular analysis revealed that, while loss of CTC1 or TEN1 leads to defective C-strand fill-in, only CTC1 KO exhibited excessive G-OH lengthening. Here, we characterized conditional STN1 KO cells and determined that STN1 KO leads to proliferation defects and telomeric damage signaling. Moreover, STN1 KO caused genome instability in the form of anaphase bridges and micronuclei. Interestingly, these phenotypes and growth inhibition were largely dependent on telomerase activity. Our findings indicate that STN1 KO closely resembles CTC1 versus TEN1 KO and that excessive G-OH extension underlies the genome instability caused by STN1 deletion.

**SUMMARY STATEMENT:** The STN1 subunit of the single-stranded DNA binding protein CST prevents telomeric damage signaling, genome instability, and proliferation defects by limiting telomerase activity.

## INTRODUCTION

Human CTC1-STN1-TEN1 (CST) is a single-stranded DNA (ssDNA) binding protein that functions in various genome maintenance pathways, including telomere length regulation, DNA replication, and DNA repair (Lim and Cech, 2021; Lyu et al., 2021b; Olson and Wuttke, 2024; Stewart et al., 2018). CST consists of numerous oligonucleotide-oligosaccharide binding folds (OB-folds), which allow it to bind to ssDNA (Lim et al., 2020). It shares homology with the well-characterized ssDNA binding protein RPA (Barbour and Wuttke, 2023; Price et al., 2010). Like RPA, the use of multiple OB-folds allows CST to bind to DNA in a length-dependent manner and to different DNA configurations (Bhattacharjee et al., 2016). When all OB-folds are engaged, it binds in the low to sub nanomolar range (Bhattacharjee et al., 2016; Chen et al., 2012). However, unlike RPA, CST has some sequence preference for G-rich DNA (Hom and Wuttke, 2017).

A primary interacting partner of CST is DNA polymerase α (pol α)-primase, an interaction conserved from yeast to humans (Giraud-Panis et al., 2010; Price et al., 2010). CTC1 and STN1 were originally discovered in a screen for factors that co-purified with pol α-primase (Goulian et al., 1990). CST stimulates the primase to polymerase transition as well as both primase and polymerase activities of pol α-primase (Casteel et al., 2009; Ganduri and Lue, 2017; Goulian and Heard, 1990; Goulian et al., 1990). This interaction is critical for CST function in telomere replication as well as DNA repair (Cai and de Lange, 2023; Mirman et al., 2022). In addition to pol α-primase, CST also interacts with other proteins involved in DNA replication and repair, such as the MCM2-7 complex, cohesin, shieldin, RAD51, MRE11, RECQ4, OGG1, and DNA polymerase β (Chastain et al., 2016; Li et al., 2023; Lyu et al., 2021a; Mirman et al., 2018; Schuck et al., 2021; Wang et al., 2019; Wysong et al., 2024). These common but varied interactions suggest that CST acts as a general genome stability factor. The extent of its involvement in each pathway is still under investigation; however, telomere maintenance is thought to be its primary function.

Telomeres consist of repetitive DNA sequences (TTAGGG in humans) at the ends of chromosomes (Shay and Wright, 2019). In humans, these DNA regions can be several to tens of kilobases in length and consist of both a double-stranded and short ssDNA (100-300 nucleotide) region. The G-rich ssDNA region, located at chromosome termini, is known as the G-overhang (G-OH). Telomeres are bound by a protection complex called shelterin, which blocks the ends from being recognized as DNA breaks and degradation by nucleases (de Lange, 2018).

Along with shelterin, CST functions in telomere length regulation. Telomere replication occurs in three distinct steps (Bonnell et al., 2021; Stewart et al., 2012a). First, the telomere duplex region is replicated by the replisome with the help of additional accessory factors, including CST (Brenner and Nandakumar, 2022; Stewart et al., 2012b). In telomerase-positive cells (e.g., germline, stem, and most cancer cells), the G-OH is then extended through the reverse transcriptase activity of telomerase (Greider and Blackburn, 1985; Morin, 1989). Following initial extension, CST binds to the G-OH to inhibit telomerase from rebinding and overextending the G-OH (Chen et al., 2012; Zaug et al., 2021). Most of the G-OH is then converted to duplex DNA by pol α-primase, leaving a short overhang for formation of the protective telomeric-loop (Olson et al., 2022). The fill-in process, known as C-strand fill-in, is facilitated by CST-dependent stimulation of pol α-primase activity (Takai et al., 2024; Wang et al., 2012; Zaug et al., 2022). Recent work also demonstrated that the shelterin subunit POT1, which protects the G-OH, recruits CST to regulate telomerase inhibition and C-strand fill-in (Cai et al., 2024).

To better understand the cellular roles of human CST, cell lines that conditionally deleted either CTC1 or TEN1 were previously generated (Feng et al., 2018; Feng et al., 2017). Conditional KO of CTC1 results in significant growth defects and the accumulation of G2 cells starting around ten days after CTC1 deletion (Ackerson et al., 2020; Feng et al., 2017). Molecular analysis of these cells demonstrated a severe defect in telomere length regulation, with G-OHs reaching kilobases in length and telomeric damage signaling (i.e., RPA and γH2AX localization). Additional analysis revealed that the elongated G-OHs were caused by the inability to inhibit telomerase and stimulate C-strand fill-in. Once overextended, there is insufficient POT1 to bind and protect the G-OH. Typically, POT1 protects G-OHs from repair, but due to the insufficient level of POT1, the now exposed ssDNA is bound by RPA, which leads to damage signaling mediated by the DNA damage response kinase ATR (Denchi and de Lange, 2007; Glousker et al., 2020). In contrast to CTC1 KO, conditional TEN1 KO exhibited no significant growth defect or telomeric damage signaling (Feng et al., 2018). Instead, there was a minor increase in G-OH length, similar to what was previously observed with shRNA knockdown of each CST subunit (Kasbek et al., 2013; Miyake et al., 2009; Surovtseva et al., 2009; Wang et al., 2012). Molecular characterization of the KO cells showed that CTC1-STN1 can inhibit telomerase without TEN1, but TEN1 is required for the stimulation of pol α-primase to complete C-strand fill-in (Feng et al., 2018).

Unlike CTC1 and TEN1, a STN1 KO cell line has not yet been characterized. The goal of this study was to characterize conditional STN1 KO cells and compare whether the phenotype resembles CTC1 or TEN1 KO, or an intermediate phenotype. We determined that deletion of STN1 closely resembles conditional deletion of CTC1, causing proliferation defects, telomeric RPA binding, and telomeric damage signaling. Moreover, we determined that STN1 KO increased general genome instability, in the form of anaphase bridges and micronuclei. To test whether these phenotypes were related to excessive G-OH lengthening, we treated STN1 KO cells with a telomerase inhibitor. Interestingly, we found that anaphase bridges, micronuclei, and cell growth were almost completely rescued following telomerase inhibition. These findings suggest that STN1 is necessary to inhibit telomerase to prevent excessive G-OH extension, genome instability, and growth inhibition.

## RESULTS

### STN1 deletion leads to decreased proliferation and G2 arrest

To assess the phenotypes associated with complete loss of STN1, we generated an inducible STN1 KO cell line. Single-guide RNAs to STN1 (sgSTN1) were generated, cloned into a retroviral vector, and transduced into previously characterized HeLa iCas9 cells, which contain Cas9 under an inducible doxycycline (DOX) promoter (McKinley, 2018; McKinley and Cheeseman, 2017). DOX was then added, and individual clones screened for loss of STN1. One of the clones that displayed efficient gene disruption, which had little to no detectable STN1 after 4 days was then selected for further characterization (Fig. 1A and S1A).

**Figure 1.**
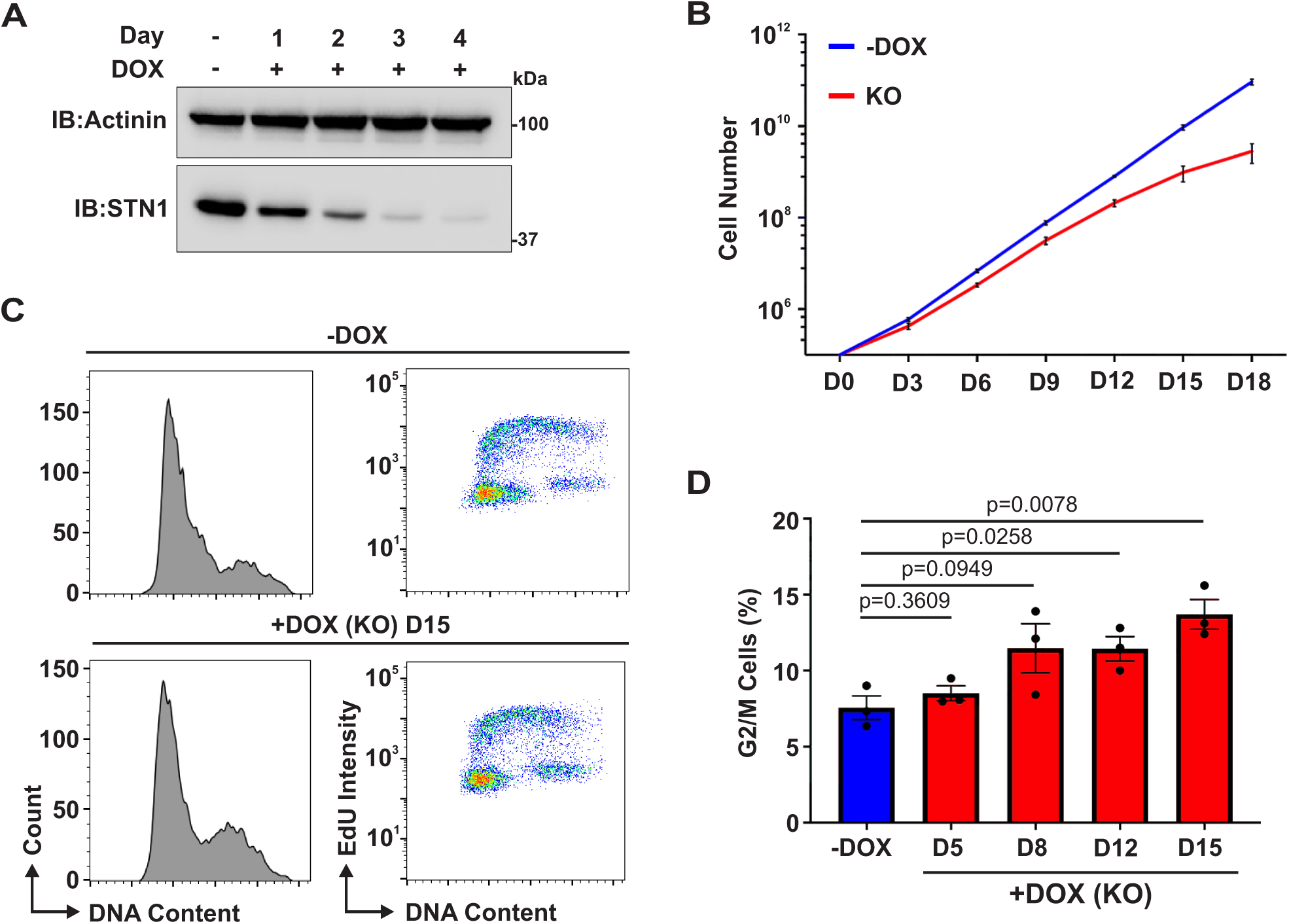
STN1 KO leads to defects in cell proliferation. (A) Western blot of STN1 levels in HeLa iCas9 sgSTN1 cells, as indicated. Actinin serves as a loading control. DOX=Doxycycline. (B) Growth curve analysis of STN1 KO and control cells. D=day after DOX addition. n=3 independent, biological replicates. (C-D) Flow cytometry analysis of STN1 KO cells. (C) Representative histograms of STN1 KO and control (-DOX) cells. Left, histograms of DNA content versus cell count. Right, DNA content versus EdU intensity. (D) Percentage of G2/M cells over time in STN1 KO cells. n=3 independent, biological replicates. Average values in the graphs indicate the mean and error bars denote ±s.e.m. *P*-values were calculated by a two-tailed, unpaired *t*-test.

Growth curve analysis demonstrated our STN1 KO cells had a significant growth defect starting around day 9 after gene disruption (Fig. 1B). The timing of this growth defect is comparable to that observed in CTC1 KO cells (Feng et al., 2017). Extension of the growth curve to day 30 showed that growth was restored after ∼21 days (Fig. S1B). Western blot analysis of STN1 from day 5 to day 25 showed little to no detectable STN1 until around day 15, after which STN1 levels began to steadily increase (Fig. S1C). These results suggest that STN1 is not deleted in a small subset of the sgSTN1 cells following DOX treatment, and that these cells eventually outgrow the KO cells to become the dominant population. This finding is not unexpected in a conditional CRISPR-Cas9 KO cell line, where there could be incomplete editing and/or breaks that are repaired in-frame in a small subset of cells. The absence of STN1 gene disruption in cells could also arise from a lack of Cas9 expression or the absence of sgSTN1. Due to the outgrowth of cells expressing STN1 starting around day 18 and rescue of cell proliferation around day 21, we confined our studies to days 5-15 when there is little to no detectable STN1 and growth is significantly decreased.

Previously, we determined that following CTC1 KO, there was an accumulation of G2-phase cells (Ackerson et al., 2020). Therefore, we performed cell cycle analysis between days 5-15 (Fig. 1C-D and S1D). This analysis showed an increase in G2/M cells starting around day 8. To distinguish between G2- and M-phase, the mitotic index, as measured by phosphorylated histone H3 S10, was then determined (Fig. S1E). The percentage of mitotic cells was unaffected in the STN1 KO compared to control cells. These results indicate that deletion of STN1 leads to the accumulation of G2 cells, similar to CTC1 KO (Ackerson et al., 2020). Combined, these results indicate that STN1 KO leads to defects in cell proliferation and cell cycle progression.

### Loss of STN1 results in telomeric RPA and damage signaling

The accumulation of G2 cells and proliferation defect did not occur until around 9 days after DOX addition, suggesting that, like CTC1 KO, these phenotypes may arise from RPA binding and ATR-mediated telomeric damage signaling due to G-OH overextension (Ackerson et al., 2020; Feng et al., 2017). To confirm this, we measured the number of RPA foci in STN1 KO cells between days 5-15 by immunofluorescence (IF) (Fig. 2A-B). We found that the number of cells with RPA foci increased significantly over time, reaching a peak at day 12. To confirm that the RPA foci were telomeric, IF-fluorescence *in situ* hybridization (FISH) analysis was then performed with an antibody to RPA and a telomere probe (Fig. S2). As expected, the vast majority of the RPA foci were localized to telomeres, similar to CTC1 KO (Ackerson et al., 2020).

**Figure 2.**
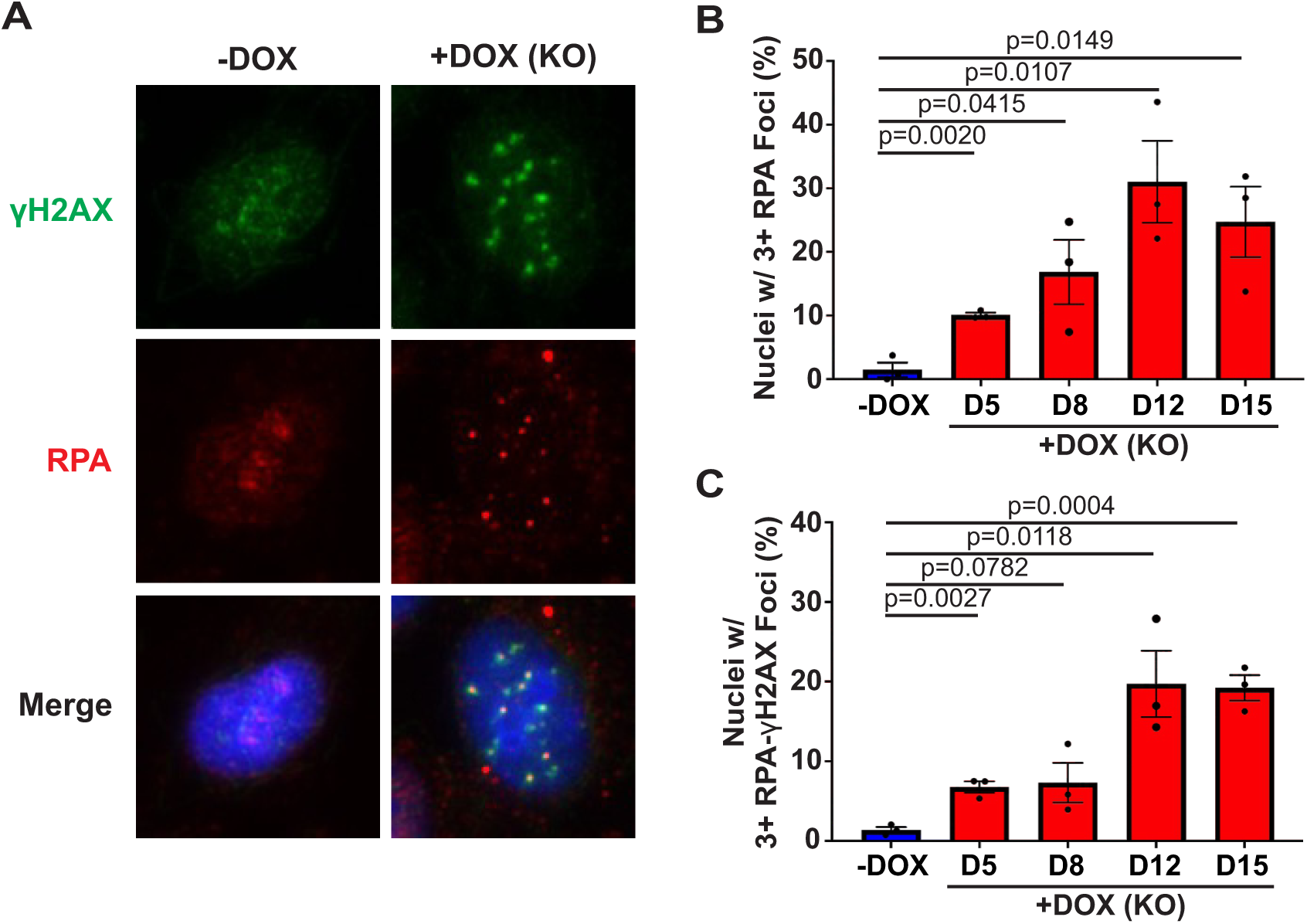
STN1 KO causes RPA-γH2AX foci. (A) Representative image of RPA-γH2AX foci. STN1 KO image is from day 12 after DOX addition. (B-C) Analysis of foci in STN1 KO and control (-DOX) cells. Percentage of nuclei displaying 3 or more RPA (B) or RPA-γH2AX foci (C). n=3 independent, biological replicates. Average values in the graphs indicate the mean and error bars denote ±s.e.m. *P*-values were calculated by a two-tailed, unpaired *t*-test. DOX=Doxycycline. D=day after DOX addition.

Next, we determined whether RPA-binding resulted in telomeric damage signaling. Co-localization between RPA and γH2AX was measured in STN1 KO cells between day 5 and day 15 (Fig. 2A and C). While there was an increase in cells with RPA-γH2AX foci on days 5 and 8, the levels more than doubled at days 12 and 15. These days correspond to the decreased proliferation and increased G2-phase cells after STN1 KO (see Figs. 1 and S1). Combined, these studies indicate that STN1 KO leads to similar defects in telomere length regulation, leading to RPA binding and telomeric damage signaling, as was observed with CTC1 KO (Ackerson et al., 2020; Feng et al., 2017).

### Add back of STN1 rescues growth and RPA foci in STN1 KO cells

To validate that the phenotypes observed were due to STN1 deletion and not off-target defects, we created a cell line expressing an sgRNA-resistant Flag-STN1 (KO+Flag-STN1). We confirmed the expression of Flag-STN1 by Western blot (Fig. 3A). The number of cells with RPA foci was then measured on day 15, which corresponds to severe growth inhibition and telomeric damage signaling (see Figs. 1-2). The expression of Flag-STN1 rescued the RPA foci (Fig. 3B). Growth curve and cell cycle analysis were then performed, and again, expression of Flag-STN1 rescued both cell growth and the increase in G2-phase cells (Fig. 3C-D), indicating these phenotypes result from STN1 deletion and not potential off-target defects of the CRISPR-Cas9 system.

**Figure 3.**
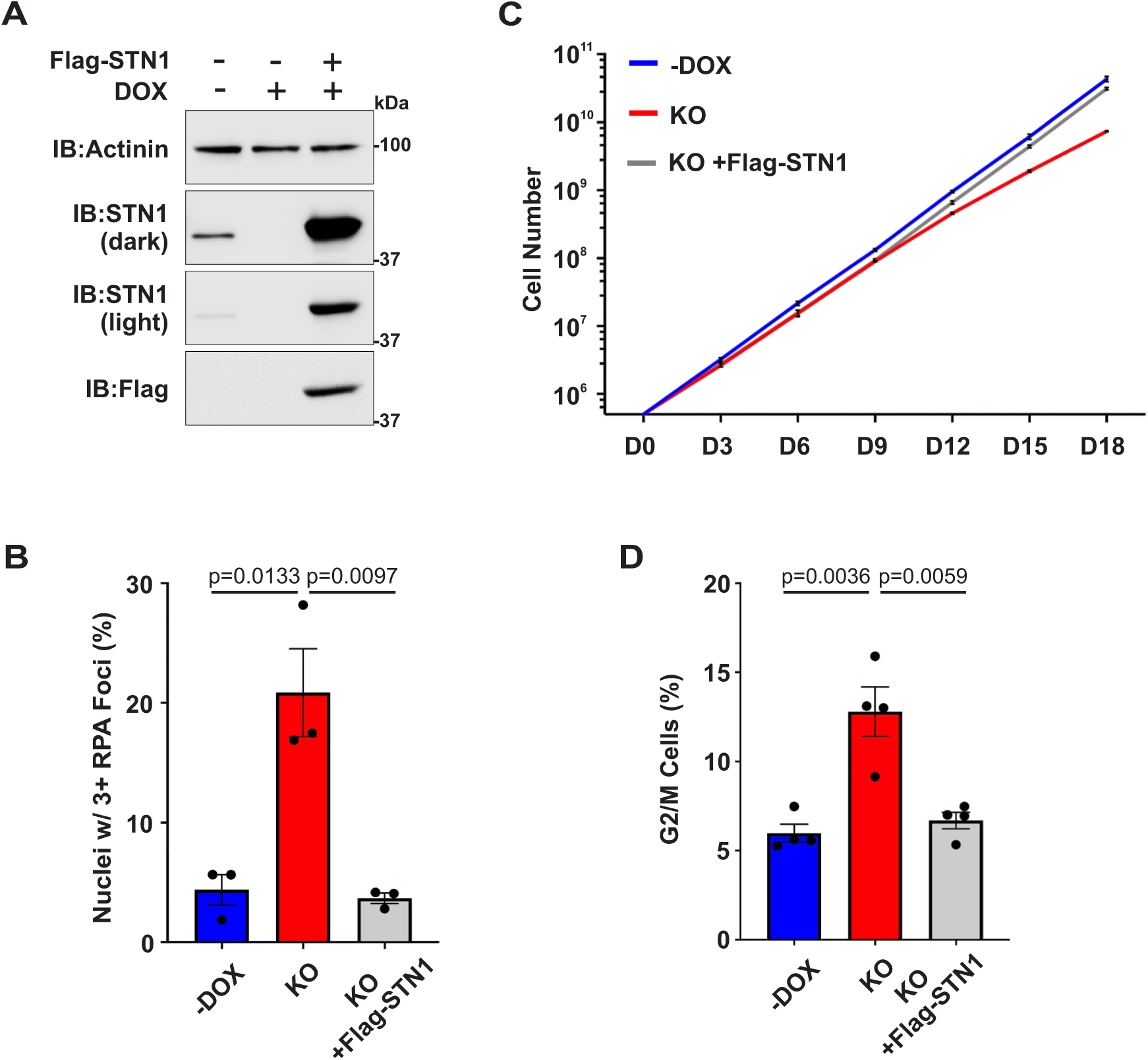
Expression of Flag-STN1 recuses RPA foci and cell proliferation defects in STN1 KO cells. (A) Western blot of STN1 and Flag-STN1 levels, as indicated. KO+Flag-STN1 denotes STN1 KO cells expressing Flag-STN1. Actinin serves as a loading control. (B) Percentage of nuclei displaying 3 or more RPA foci on day 15 after DOX addition. n=3 independent, biological replicates. (C) Growth curve analysis, as indicated. n=3 independent, biological replicates. (D) Percentage of G2/M cells determined by flow cytometry on day 15 after DOX addition. n=4 independent, biological replicates. Average values in the graphs indicate the mean and error bars denote ±s.e.m. *P*-values were calculated by a two-tailed, unpaired *t*-test. DOX=Doxycycline.

### STN1 deletion causes general genome instability phenotypes

We and others previously determined that STN1 knockdown with shRNA or siRNA increases general genome instability in the form of anaphase bridges and micronuclei (Bhattacharjee et al., 2016; Lyu et al., 2021a; Stewart et al., 2012b; Surovtseva et al., 2009). To determine whether STN1 KO also displayed these phenotypes, we measured the levels of anaphase bridges and micronuclei between days 5-15 (Fig. 4). There was a significant increase in both phenotypes starting at day 5, with the levels continuing to increase through day 15 (Fig. 4B and E). Expression of Flag-STN1 rescued the increases in both anaphase bridges and micronuclei on day 15 (Fig. 4C and F). These findings indicate that STN1 KO results in genome instability.

**Figure 4.**
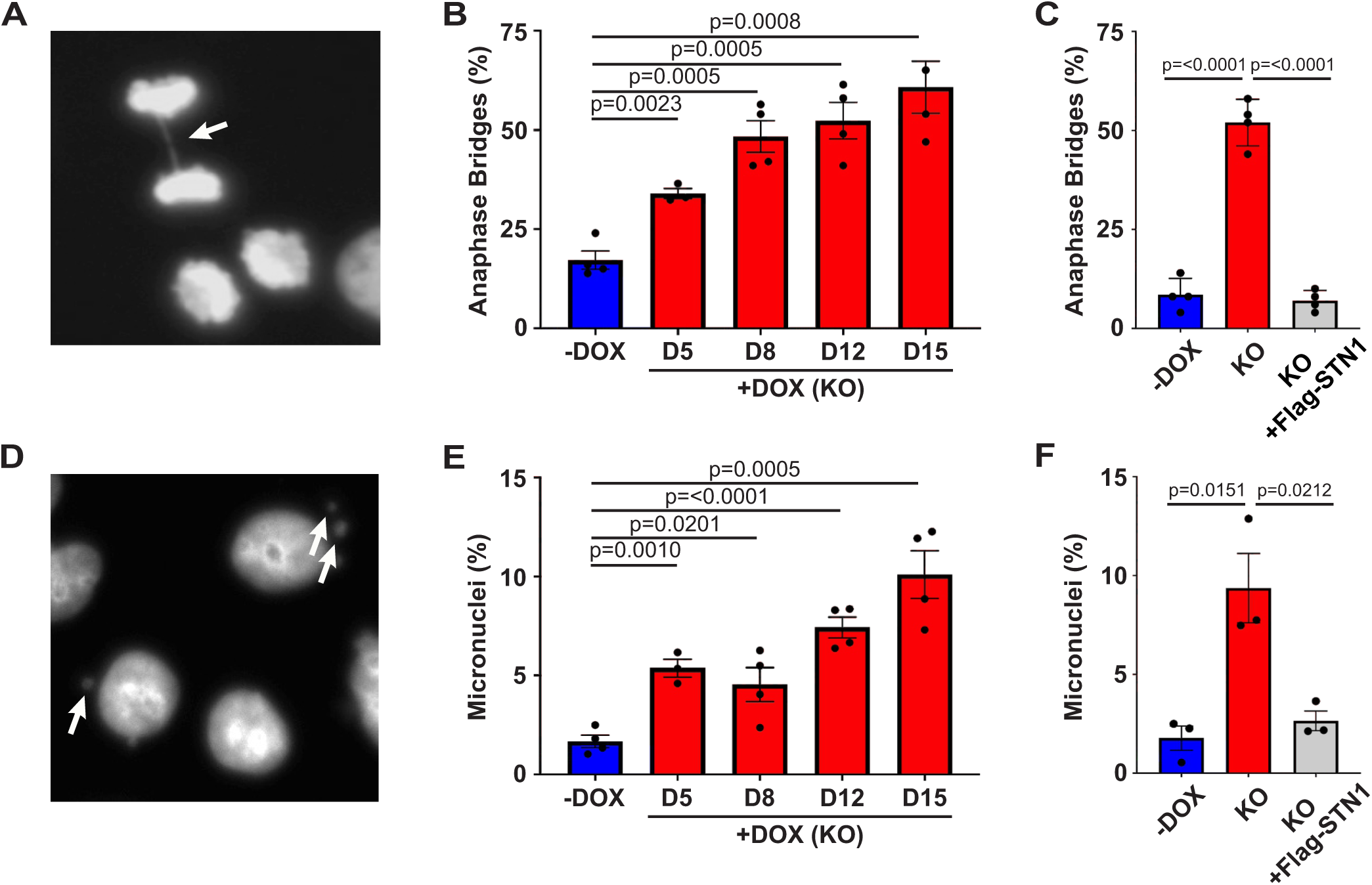
STN1 KO increases general genome instability. (A) Representative image of an anaphase bridge (indicated by the arrow) in STN1 KO cells. (B-C) Percentage of anaphase bridges in STN1 KO and control (-DOX) cells between days 5-15 (B) or with expression of exogenous Flag-STN1 on day 15 (C). n=≥3 independent, biological replicates. (D) Representative image of micronuclei (indicated by the arrows) in the STN1 KO cells. (E-F) Percentage of cells with micronuclei in STN1 KO and control cells between days 5-15 (E) or with expression of exogenous Flag-STN1 on day 15 (F). n=≥3 independent, biological replicates. Average values in the graphs indicate the mean and error bars denote ±s.e.m. *P*-values were calculated by a two-tailed, unpaired *t*-test. DOX=Doxycycline. D=day after DOX addition.

### Inhibition of telomerase largely rescues genome instability and cell growth

CST is known to play distinct roles in telomere maintenance, DNA replication, and DNA repair. However, how loss of CST function in these different processes affects genome stability is currently unknown. Therefore, we determined to test whether the anaphase bridges and micronuclei in the STN1 KO cells correlated with G-OH elongation and subsequent telomeric damage signaling. For this analysis, cells were treated with the well-characterized telomerase inhibitor BIBR1532 (which will be referred to simply as BIBR) (Damm et al., 2001; Pascolo et al., 2002). We treated the cells with 20 μM BIBR or DMSO starting on day 0. Treatment continued until fixation of the cells for IF on day 15. Previous work from the Cech lab showed that treatment with 20 μM BIBR significantly inhibited telomerase in HeLa cells (Nakashima et al., 2013). BIBR treatment was also shown to limit G-OH overextension in CTC1 KO cells (Feng et al., 2018).

Following treatment with BIBR, we observed a significant rescue in cells with RPA and RPA-γH2AX foci, indicating a reduction in G-OH overextension and telomeric damage signaling (Fig. 5A-B). We next measured the levels of anaphase bridges and micronuclei on day 15 following STN1 KO with and without BIBR-treatment. Surprisingly, BIBR treatment almost completely rescued both the increased anaphase bridges and micronuclei (Fig. 5C-D), suggesting that these phenotypes are mainly dependent on excessive G-OH elongation by telomerase and not non-telomere phenotypes. However, we do note that the anaphase bridges and micronuclei were not completely rescued following BIBR treatment, suggesting that some portion of the anaphase bridges and micronuclei may be independent of G-OH elongation and telomere damage signaling (see Discussion for more details).

**Figure 5.**
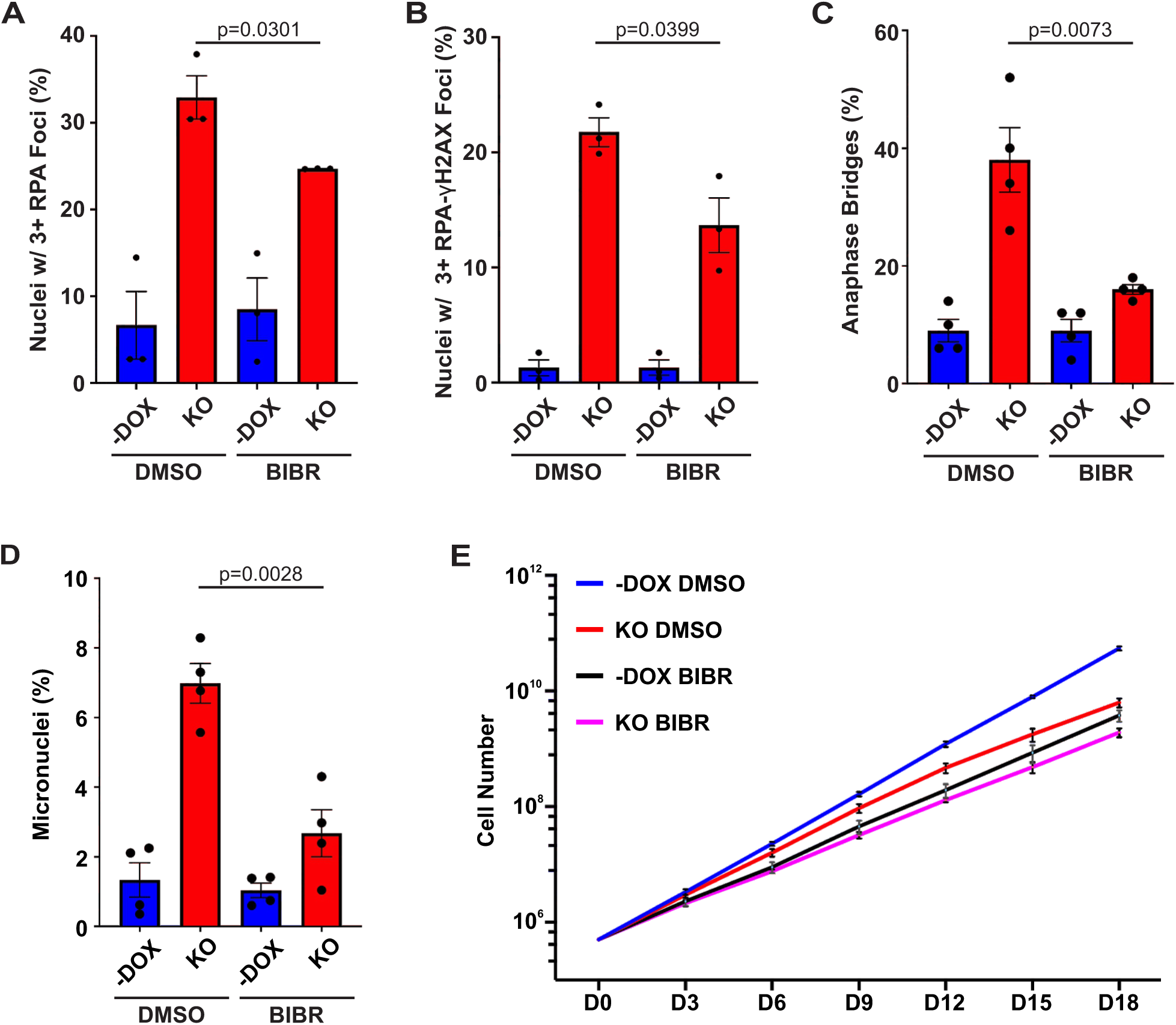
Telomerase inhibition largely rescues anaphase bridges, micronuclei, and cell growth. Cells were treated with a telomerase inhibitor (BIBR) or DMSO starting with the addition of DOX on day 0 to the sgSTN1 cells. The control cells were also treated with BIBR or DMSO across the same timeframe. (A-B) Percentage of nuclei with 3 or more RPA (A) or RPA-γH2AX (B) foci in STN1 KO or control cells with or without BIBR treatment. n=3 independent, biological replicates. (C-D) Percentage of micronuclei (C) or anaphase bridges (D) on day 15 after DOX addition with and without BIBR treatment. n=4 independent, biological replicates. (E) Growth curve analysis, as indicated. n=3 independent, biological replicates. Average values in the graphs indicate the mean and error bars denote ±s.e.m. *P*-values were calculated by a two-tailed, unpaired *t*-test. DOX=Doxycycline. D=day after DOX addition.

The results described above posit that the growth defect could also be primarily caused by telomere dysfunction since genome instability often precedes cell death. Consequently, we tested how cell growth was affected following telomerase inhibition (Fig. 5E). The previously mentioned study from the Cech lab observed a growth defect with BIBR-treatment (Nakashima et al., 2013). We found a similar effect in the HeLa iCas9 cells (Fig. 5E, compare blue to black line). Comparison of the STN1 KO and - DOX cells after DMSO treatment showed a similar defect in cell proliferation following STN1 deletion (Fig. 5E, compare red to blue line). However, comparison of STN1 KO to -DOX cells treated with BIBR showed only a minor decrease (Fig. 5E, compare purple to black line). Comparison of the total number of cells at day 18 confirm an almost 10-fold decrease in the DMSO-treated STN1 KO cells compared to the control (-DOX DMSO: 5.5×10^10^, KO DMSO: 6.3×10^9^), whereas there was a less than 2-fold decrease with the BIBR-treated cells (-DOX BIBR: 3.7×10^9^ and KO BIBR: 1.9×10^9^).

These findings suggest that telomerase inhibition almost completely rescues cell proliferation in the STN1 KO cells. The minor decrease observed in the BIBR-treated KO cells is likely due to the loss of non-telomere CST functions or incomplete rescue of telomeric damage signaling (Fig. 5A-B). Regardless, these results indicate that genome instability and defects in cell proliferation arise from an inability of CST to inhibit telomerase, which results in G-OH overextension and telomeric damage signaling, versus non-telomere defects associated with CST loss.

## DISCUSSION

Previous work considered the effects of CTC1 or TEN1 KO in human cells (Ackerson et al., 2020; Feng et al., 2018; Feng et al., 2017). In this work, we studied the impact of conditional STN1 KO. Loss of STN1 resulted in severe growth defects, the accumulation of G2-phase cells, and telomeric damage signaling from excessive G-OH elongation. These findings suggest that STN1 KO resembles that of CTC1 versus TEN1 KO. Moreover, we show that STN1 KO leads to significant increases in anaphase bridges and micronuclei. These phenotypes, along with the defects in cell proliferation, are largely dependent on telomerase activity, suggesting that the inability to prevent telomerase from overextending the G-OH, versus other genome maintenance functions of CST, is a major cause of genome instability in unperturbed conditions.

### Functional roles of CST subunits in telomere replication

CTC1 and TEN1 KO studies were performed in HCT116 cell lines by inserting loxP sites and then inducing Cre nuclease nuclear localization to remove the floxed region (Feng et al., 2018; Feng et al., 2017). Efforts to create a similar cell line for STN1 were unsuccessful, so we used a previously characterized HeLa inducible Cas9 cell line to conditionally delete STN1, using CRISPR-Cas9 technology (Cong et al., 2013; Jinek et al., 2013). Even though the cell type and mode of gene disruption differ between the CTC1 and STN1 KO lines, the phenotypes closely mirror one another. Deletion of CTC1 or STN1 has a profound, but not immediate, impact on cell growth and cell cycle progression (Ackerson et al., 2020; Feng et al., 2017) (Fig. 1). In contrast, deletion of TEN1 has little impact on cell proliferation (Feng et al., 2018).

Biochemical reconstitution demonstrates how CST orchestrates telomere replication by limiting telomerase activity and then stimulating pol α-primase activity to complete C-strand fill-in (Takai et al., 2024; Zaug et al., 2022; Zaug et al., 2021). The contributions of each subunit to these essential processes are highlighted by these gene KO experiments (Ackerson et al., 2020; Feng et al., 2018; Feng et al., 2017) (this study). Loss of either CTC1 or STN1 leads to excessive, telomerase-dependent G-OH lengthening, leading to telomeric RPA and γH2AX, whereas TEN1 KO leads to a modest ∼2-fold increase in G-OH signal and the absence of telomeric damage signaling (Feng et al., 2018). Analysis of CTC1 KO cells demonstrated that the overextended G-OH eventually depletes the available pools of POT1, leading to RPA binding, ATR localization, and H2AX phosphorylation (Ackerson et al., 2020; Feng et al., 2017). Consistent with this model, cells derived from CTC1 KO mice showed G-OH overextension, telomeric γH2AX foci, and cell proliferation defects (Gu et al., 2012). Furthermore, we show in this study that growth inhibition following STN1 deletion is dependent on telomerase activity (Fig. 5E).

It is interesting to note that, while telomeric RPA foci occur at earlier time points, the growth defect in CTC1 and STN1 KO cells does not occur until later, around day 9 (Fig. 1) (Feng et al., 2017). In the STN1 KO, there is a 2.7-fold increase in RPA-γH2AX foci from day 8 to day 12, which correlates with growth inhibition (Fig. 1-2). An ∼20% increase in telomeric γH2AX foci from day 7 to day 12 was also reported following CTC1 KO (Feng et al., 2017). Furthermore, reducing the levels of RPA-γH2AX with BIBR-treatment is sufficient to rescue cell growth (Fig. 5E). These results suggest that a certain level of ATR-mediated telomeric damage signaling is tolerated before inducing global cell cycle arrest. This is consistent with previous studies demonstrating that the recruitment of ATR-ATRIP and associated ATR activators to RPA-bound DNA – sufficient to induce global checkpoint activation – is length-dependent (Bantele et al., 2019; Choi et al., 2010; Cortez et al., 2001; Saldivar et al., 2017; Zou and Elledge, 2003).

We therefore propose that the growth inhibition caused by G-OH overextension requires the accumulation of sufficient telomeric RPA to activate a global, ATR-mediated checkpoint response, followed by senescence or apoptosis. (Both senescence and apoptosis have been observed in CTC1 KO cells (Ackerson et al., 2020; Feng et al., 2017; Gu et al., 2012).) This model highlights the essential roles of POT1 and CST in telomere protection, with POT1 preventing RPA association and CST, which is recruited by POT1, inhibiting telomerase to prevent G-OH elongation, which can lead to POT1 exhaustion and subsequent RPA binding. Additional work will be necessary to determine the timing of ATR-ATRIP recruitment to overextended G-OHs and how this then leads to checkpoint activation, particularly since the G2 arrest in CTC1 KO cells was ATR-dependent but CHK1-independent (Ackerson et al., 2020).

### Telomere dysfunction leads to general genome instability following CST loss

The central role of CST in telomere replication is conserved from yeast to humans (Giraud-Panis et al., 2010; Price et al., 2010). However, at least in mammals, CST plays significant roles in genome maintenance outside telomeres. CTC1 and STN1 were initially discovered through a screen for pol α-primase interacting partners (Goulian et al., 1990). Named AAF (α-accessory factor), this complex was shown to stimulate pol α-primase activity and was thought to aid in DNA replication (Goulian and Heard, 1990). Work published almost two decades later provided more direct evidence of AAF (CTC1-STN1) function in DNA replication (Casteel et al., 2009). That same year, studies were published citing the exciting discovery of CST in plants and mammals as a second telomere maintenance complex, in addition to shelterin (Miyake et al., 2009; Surovtseva et al., 2009). In *Saccharomyces cerevisiae*, a similar CST complex (Cdc13-Stn1-Ten1) plays an essential role in both telomere protection and replication (Giraud-Panis et al., 2010; Price et al., 2010). Mammalian CST functions similar to the *S. cerevisiae* CST to regulate telomere replication, whereas shelterin is the major protective complex.

Our work and that of many others have uncovered important non-telomeric functions of CST. This includes roles in DNA replication restart, double-strand break repair, origin licensing, base excision repair, protection of stalled replication forks, and maintenance of sister chromatid cohesion (Chastain et al., 2016; Hara et al., 2023; Lyu et al., 2021a; Lyu et al., 2021b; Mirman et al., 2018; Schuck et al., 2021; Stewart et al., 2012b; Wang et al., 2019; Wysong et al., 2024). Previous and more recent work indicates that *S. cerevisiae* CST also functions in DNA replication (Calvo et al., 2019; Gasparayan et al., 2022; Gasparyan et al., 2009); however, its primary role is telomeric.

Based on the previous AAF work, the diverse functions of human CST, and the evolution of the shelterin complex for telomere protection, it remained unclear whether CST’s primary role is at telomeres. In this study, we mitigated the major consequences of CST loss at telomeres by treating our STN1 KO cells with a telomerase inhibitor (Figure 5). This allowed us to test whether the growth defect and general genome instability phenotypes correlate with CST-associated telomere dysfunction and begin addressing the primary function of CST. Our initial hypothesis was that these phenotypes would be only partially rescued following BIBR-treatment since defects in cell proliferation and genome stability have been observed following treatment with exogenous DNA-damaging or fork-stalling agents (Chastain et al., 2016; Stewart et al., 2012b; Wang et al., 2014). However, the increase in anaphase bridges and micronuclei and decreased proliferation were predominantly rescued with telomerase inhibition, suggesting telomere length regulation is the primary role of human CST, and its other genome maintenance functions are more specialized.

### Potential implications for understanding CST-associated telomere biology disorders

CST mutations are associated with the telomere biology disorders Coats plus (CP) and dyskeratosis congenita (DC) (Anderson et al., 2012; Keller et al., 2012; Polvi et al., 2012; Simon et al., 2016). Telomere biology disorders are broadly characterized by abnormal telomere length (Armanios, 2022; Armanios and Blackburn, 2012; Revy et al., 2023; Savage, 2022). DC patients have germline mutations in a variety of telomere maintenance factors, which lead to severe telomere shortening, with a wide range of clinical manifestations, particularly in high-turnover tissues, and an increased risk of certain cancers. Patients often succumb to bone marrow failure. CP is a more severe form of DC with additional features, such as retinopathy, intrauterine growth restriction, intracranial calcifications, osteopenia, and gastrointestinal tract bleeding. Most genetic mutations identified in CP patients have been found in *CTC1* and *STN1* (Grill and Nandakumar, 2021).

Given the multiple functions of CST in genome maintenance and the additional features of CP, it has been unclear whether the cellular defects that give rise to CP are strictly telomeric in nature or a combination of defects in telomere and non-telomere CST functions. Interestingly, not all CP patients displayed telomere shortening, and characterization of *CTC1* and *STN1* CP mutations showed both telomeric and general genome instability phenotypes (Anderson et al., 2012; Simon et al., 2016; Walne et al., 2013; Wang and Chai, 2018). In particular, characterization of patient-derived cell lines expressing *STN1* CP mutations found decreased proliferation and increased genome instability, including micronuclei, nuclear bridges, and replication defects, but not significant changes in telomere length compared to the healthy controls (Simon et al., 2016). However, the telomeric and non-telomeric phenotypes have been mixed across the spectrum of *CTC1* and *STN1* mutations studied and do not follow a clear pattern (Chen et al., 2013; Gu and Chang, 2013; Gu et al., 2018; Wang and Chai, 2018).

Our work suggests that the growth defect, which may be associated with intrauterine growth restriction and the depletion of stem cell pools, and the general genome instability observed could result from an inability to limit telomerase activity as opposed to instability from the loss of non-telomeric CST functions in genome maintenance. However, defects in C-strand fill-in are also likely to contribute to CP since mutations in *POLA2*, a subunit of pol α-primase, were recently uncovered in two separate families with a telomere biology disorder expressing CP features (Kvarnung et al., 2025). Furthermore, treatment of cells expressing a CP *POT1* mutation with the telomerase inhibitor BIBR only partially rescued G-OH lengthening (Takai et al., 2016). Therefore, further studies will be necessary to elucidate the contributions of telomerase inhibition versus defective C-strand fill-in of CP-associated CST mutations and how this contributes to features of the disease.

## MATERIALS AND METHODS

### Cell culture

HeLa iCas9 cells were maintained in DMEM supplemented with 10% fetal bovine serum and 1% penicillin/ streptomycin and grown at 37°C with 5% CO_2_. Cell lines were regularly checked for contamination. HeLa iCas9 (inducible Cas9) cells were generously provided by Dr. Iain Cheeseman from the Massachusetts Institute of Technology (McKinley and Cheeseman, 2017). Generation of the STN1 inducible knockout (KO) cell line was previously described (Wysong et al., 2024). Cell lines expressing Flag-STN1 were made by transducing the HeLa iCas9 sgSTN1 cells with retrovirus generated from a pMIT vector encoding an sgRNA-resistant Flag-STN1 construct. Cells were then selected by flow cytometry for the expression of Thy1.1. To induce conditional gene KO of STN1, cells were incubated with 1 μg/ml Doxycycline (DOX) for 4-5 days. For experiments in Figure 5, cells were treated with BIBR1532 (telomerase inhibitor), which was added fresh every 2-3 days when cells were passaged. The telomerase inhibitor BIBR1532 (Selleckchem, S1186), which was dissolved in DMSO (dimethyl sulfoxide), was added at a final concentration of 20 μM starting on day 0. DMSO control samples were treated with an equivalent amount of DMSO at each cell passage.

### Growth curve

Cells were plated at 5×10^5^ and allowed to grow for 3 days. They were then counted and replated at 5×10^5^. This was repeated until the indicated time point, and the total number of cells was extrapolated from the cell counts to generate the graphs.

### Flow cytometry

Cells were collected and samples prepared as previously described (Ackerson et al., 2020), except that they were incubated for 1 hour with EdU prior to collection. The samples were then run on a BD FACSMelody (10,000 cells/biological replicate) and analysis performed using FlowJo 10 software.

### Antibodies

Primary: OBFC1 (STN1) (Novus, NBP2-01006), Actinin (Santa Cruz, sc17829), pH3 S10 (Cell Signaling, 9706), RPA32 (Abcam, ab16850), and γH2AX (Bethyl, A300-081A; Abcam, ab81299). Secondary: Invitrogen: anti-rabbit-HRP (32460); anti-mouse-HRP (32430); Invitrogen: goat-anti-rabbit AlexaFluor 488 (A11034), goat-anti-mouse AlexaFluor 488 (A11029), goat-anti-rabbit AlexaFluor 594 (A11037), goat-anti-mouse AlexaFluor 594 (A11032).

### Western blot

Cell pellets were suspended in lysis buffer (20 mM Tris pH 8.0, 100 mM NaCl, 1 mM MgCl_2_, 0.1% IGEPAL) with fresh protease and phosphatase inhibitors added and incubated at 4°C for 15 minutes with rotation. Samples were then centrifuged at 13,000 rpm for 7 minutes. The supernatant was then collected and protein concentration measured by BCA assay (Pierce). 30 µg of protein per well was run by SDS-PAGE and transferred to a nitrocellulose membrane (Cytiva Amersham) at 25V overnight. All membranes were checked with Ponceau S staining for transfer efficiency and total protein loading. Membranes were blocked with 1% non-fat milk in 1x phosphate-buffered saline (PBS) plus 0.1% Tween 20 (PBST) for at least 2 h. Primary antibodies were diluted in 1% non-fat milk-PBST and incubated at 4°C overnight (Antibody dilutions: STN1 1:2000, α-Actinin 1:10000, Flag 1:1000, Cas9 1:2000). The membranes were then washed 3x for 10 min each in PBST. Secondary antibodies were diluted in 1% non-fat milk-PBST for at least 2 h at room temperature (RT). After incubation, the blots were then developed with Western Lightning Plus ECL (Perkin Elmer) or ECL Prime (Cytiva Amersham) and imaged on a Bio-Rad ChemiDoc MP.

### Immunofluorescence (IF)

Cells were plated onto coverslips and allowed to grow for 24 hours to 50-70% confluency. For all antibodies except phosphorylated histone H3 S10 (pH3 S10), cells were pre-extracted with 0.1% Triton X-100 in CSK buffer (10 mM HEPES pH 7.4, 300 mM Sucrose, 100 mM NaCl, 3 mM MgCl_2_) with fresh protease and phosphatase inhibitors for 5 min at RT. Coverslips were then washed once with PBS and were fixed with 4% formaldehyde in PBS for 10 min at RT. After formaldehyde incubation, cells were rinsed once with PBS and then permeabilized with 0.5% Triton X-100 diluted in PBS for 10 min at RT. Following permeabilization, coverslips were washed with PBS and then stored 4°C in PBS until IF was performed. For the detection of pH3 Ser10, the pre-extraction step was not performed. IF was then performed as previously described (Ackerson et al., 2020). (Antibody dilutions: RPA 1:500, γH2AX 1:1000, pH3 S10 1:500). For IF combined with fluorescence *in situ* hybridization (IF-FISH), IF was performed as described above and then telomere FISH was performed with a telomeric G-strand PNA probe (AlexaFluor 488-[CCCTAA]_3_; PNA Bio), as previously described (Ackerson et al., 2020). Coverslips were mounted onto slides with Fluoromount-G containing 0.2 μg/ml DAPI. IF and IF-FISH images were then taken under 40x or 63x objectives, respectively, on a Leica DMi8 fluorescence microscope. Foci and co-localizing foci were determined using ImageJ. DAPI images from the IF experiments were used to determine the number of micronuclei. At least 5 images and a minimum of 150 nuclei were analyzed per condition per independent, biological replicate.

### Anaphase bridges

Cells were plated on coverslips and allowed to grow overnight. Nocodazole was then added for 2 hours, cells were then washed three times with PBS and allowed to recover for 100-120 minutes. Cells were then fixed with 3% formaldehyde in PBS for 10 min. Coverslips were washed three times with PBS and mounted onto slides with Fluoromount-G containing 0.2 μg/ml DAPI. 50 anaphase bridges were scored for each independent, biological replicate.

### Data Analysis

At least three independent, biological replicates were completed for each experimental condition, as indicated in the figure legends. *P-*values were calculated using a two-tailed, unpaired *t*-test, using GraphPad Prism 10 software.

## Acknowledgements

We would like to thank Yilin Wang for her assistance generating the HeLa iCas9 sgSTN1 cell line, Daniel Burnett, Inyeneabasi Ekrikpo, and Kennedy Vogler for their assistance with IF experiments, and members of the Stewart lab for useful discussions and critical reading of the manuscript. We would also like to thank Dr. Ian Cheeseman for generously providing the HeLa iCas9 cells and the Western Kentucky University Biotechnology Center for use of instrumentation.

## Competing interests

The authors declare no competing interests.

## Funding

This work was funded by a Research Project Award and Start-up Award from KY-INBRE (National Institutes of Health P20GM103436) to J.A.S., startup funds from the Western Kentucky University Office of Research and Creative Activity and Ogden College of Science and Engineering to J.A.S, and Faculty Undergraduate Student Engagement grants from Western Kentucky University to C.A.S, S.D.R, and G.H.D.

## Data and resource availability

All relevant data can be found within the article and its supplementary information.

## Notes

### Competing Interest Statement

The authors have declared no competing interest.

