## Supplemental Files for "Conditional deletion of human STN1 leads to telomere dysfunction and telomerase-dependent genome instability and proliferation defects"

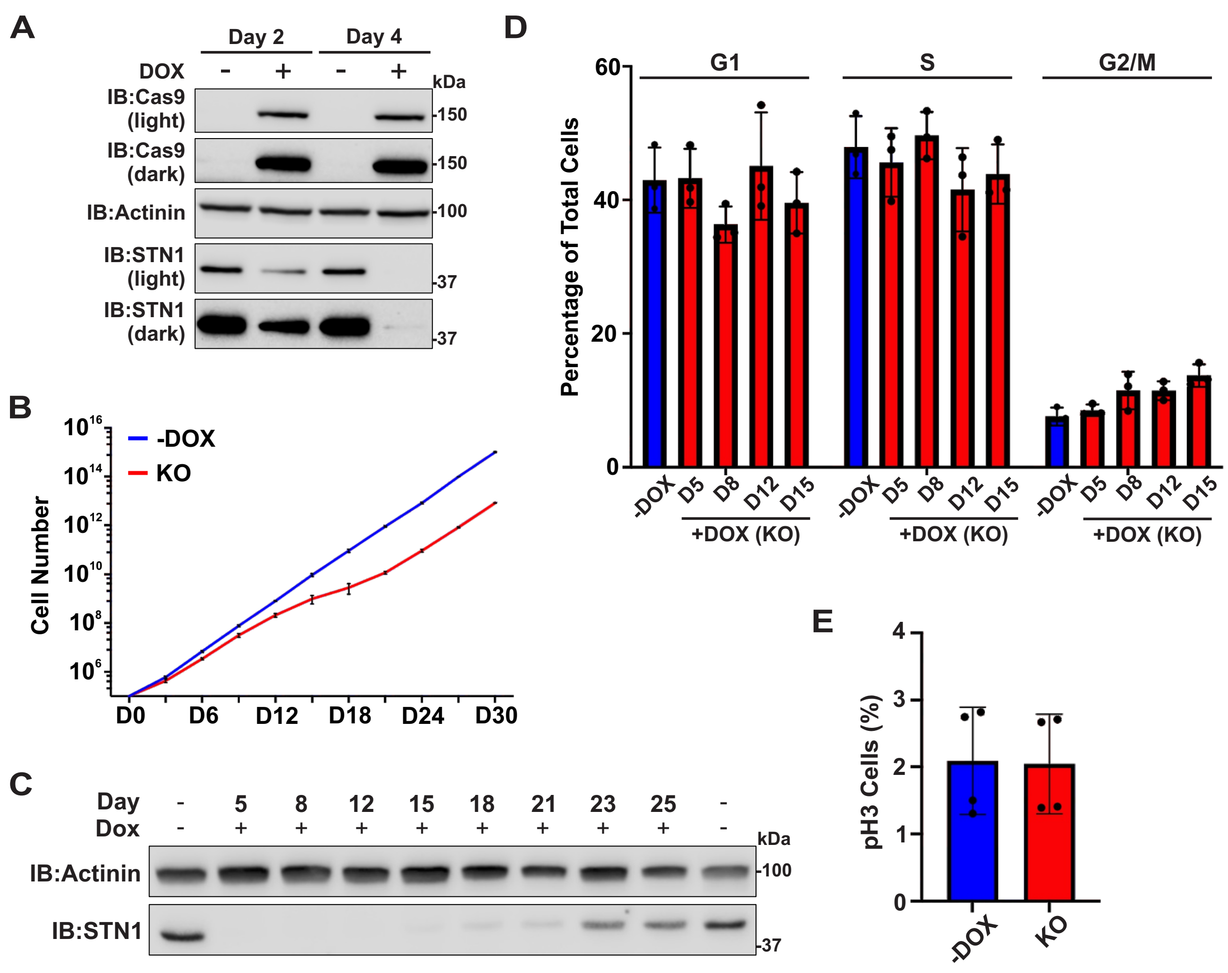


**Figure S1.** (A) Western blot of Cas9 and STN1 levels in HeLa iCas9 sgSTN1 cells, as indicated. Actinin serves as a loading control. DOX=Doxycycline. (B) Growth curve analysis of STN1 KO and control cells. D=day after DOX addition. (C) Western blot of STN1 levels over time after DOX addition. Actinin serves as a loading control. (D) Flow cytometry analysis of STN1 KO cells. Percentage of cells in different phases of the cell cycle. n=3 independent, biological replicates. (E) Percentage of mitotic cells as measured by the immunofluorescence of phosphorylated H3 S10 (pH3). n=4 independent, biological replicates. Average values in the graphs indicate the mean and error bars denote ±s.e.m. *P*-values were calculated by a two-tailed, unpaired *t*-test.

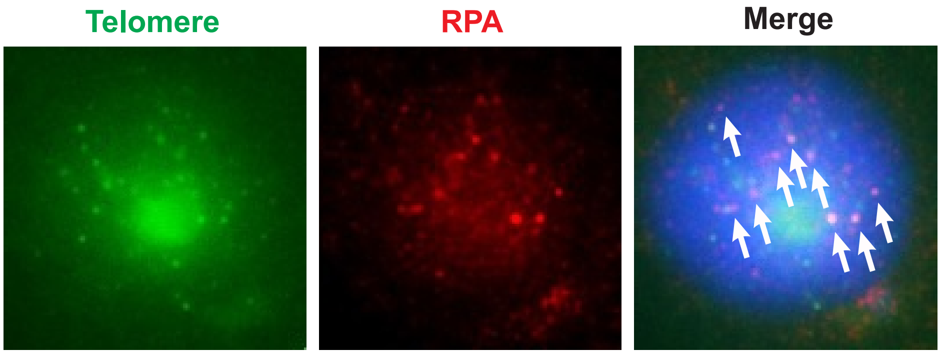


**Figure S2.** Representative image of RPA localization to telomeres in STN1 KO cells on day 15. The arrows denote co-localizations. Blue in the merged image is DAPI staining.

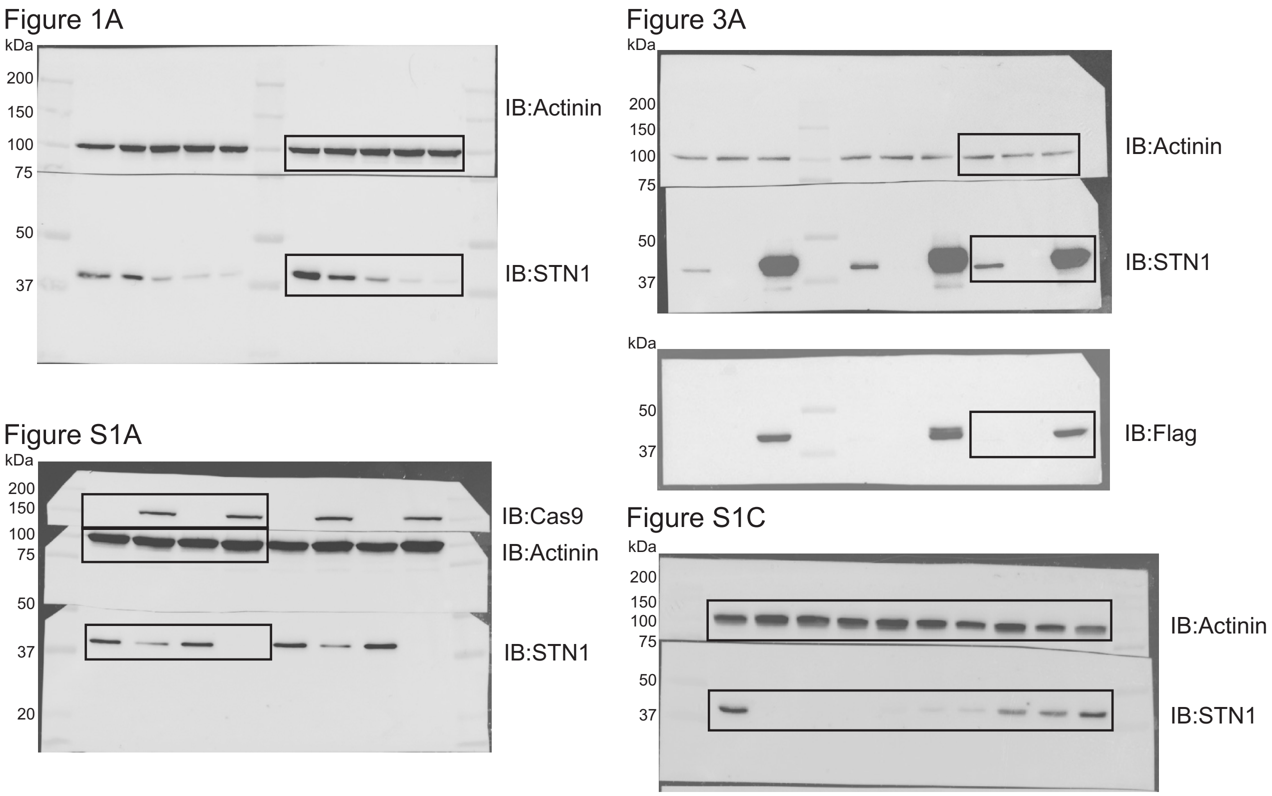


**Figure S3.** Blot transparency. Full Western blots (overlay of chemiluminescence/colorimetric images) are shown for the corresponding figure, as indicated above the blots. Boxes identify regions that were included in the figures. For Figure 3A, the STN1 blot was stripped and re-probed with an anti-Flag antibody.
